# Galectin-3-binding protein inhibits extracellular heparan 6-*O*-endosulfatse Sulf-2

**DOI:** 10.1101/2023.12.20.572603

**Authors:** Aswini Panigrahi, Julius Benicky, Reem Aljuhani, Pritha Mukherjee, Zora Nováková, Cyril Bařinka, Radoslav Goldman

**Author notes:** Corresponding author: Aswini Panigrahi.

## Abstract

Human extracellular 6-*O*-endosulfatases Sulf-1 and Sulf-2 are the only enzymes that post-synthetically alter the 6-*O* sulfation of heparan sulfate proteoglycans (HSPG), which regulates interactions of HSPG with many proteins. Oncogenicity of Sulf-2 in different cancers has been documented and we have shown that Sulf-2 is associated with poor survival outcomes in head and neck squamous cell carcinoma (HNSCC). In spite of its importance, limited information is available on direct protein-protein interactions of the Sulf-2 protein in the tumor microenvironment. In this study, we used monoclonal antibody (mAb) affinity purification and mass spectrometry to identify galectin-3-binding protein (LG3BP) as a highly specific binding partner of Sulf-2 in the secretome of HNSCC cell lines. We validated their direct interaction *in vitro* using recombinant proteins and have shown that the chondroitin sulfate (CS) covalently bound to the Sulf-2 influences the binding to LG3BP. We confirmed importance of the CS chain for the interaction by generating a mutant Sulf-2 protein that lacks the CS. Importantly, we have shown that the LG3BP inhibits Sulf-2 activity *in vitro* in a concentration dependent manner. As a consequence, the addition of LG3BP to a spheroid cell culture inhibited invasion of the HNSCC cells into Matrigel. Thus, Sulf-2 interaction with LG3BP has functional relevance, and may regulate physiological activity of the Sulf-2 enzyme as well as its activity in the tumor microenvironment.

## Introduction

Human extracellular 6-*O*-endosulfatases Sulf-1 and Sulf-2 are homologous enzymes that exhibit arylsulfatase activity and highly specific endoglucosamine-6-sulfatase activity (Dhoot et al., 2001; Morimoto-Tomita et al., 2002; Seffouh et al., 2019). These two are the only enzymes that post-synthetically alter the 6-*O* sulfation of heparan sulfate proteoglycans (HSPG) which affects binding to multiple ligands including VEGF, FGF, HGF, fibronectin, or certain chemokines (Feyzi et al., 1997; Lyon et al., 1994; Mahalingam et al., 2007; Ono et al., 1999; Pye et al., 2000; Uchimura et al., 2006; Zhang et al., 2012). These interactions regulate many impactful signaling events and fundamentally modulate various developmental processes or pathological activities in the tumor microenvironment. Studies from our and other laboratories have shown upregulation of Sulf-2 in various cancer types and its impact on patient survival (Bret et al., 2011; Flowers et al., 2016; Hur et al., 2012; Lai et al., 2008, p. 20; Phillips et al., 2012; Yang et al., 2022, 2021; Zhu et al., 2016). Thus, the Sulf-2 enzyme is expected to serve as an essential regulatory elements in HS-dependent developmental and pathophysiological processes (Rosen and Lemjabbar-Alaoui, 2010; Vivès et al., 2014).

The Sulf-2 is a secretory enzyme that have been characterized by us (Benicky et al., 2023) and others (Dierks et al., 2003; Uchimura et al., 2006; Vivès et al., 2014). The enzyme is extensively N-glycosylated (Benicky et al., 2023) and processed by furin-cleavage to a mature form consisting of a disulfide-bound pair of subunits (Morimoto-Tomita et al., 2002). The highly conserved N-terminal subunit contains the catalytic active site carrying a formylglycine residue generated from a cysteine by a dedicated SUMF1 processing enzyme (Appel and Bertozzi, 2015; Hanson et al., 2004; Tang and Rosen, 2009). Recent studies show that the C-terminal hydrophilic domain of Sulf-2 carries a chondroitin sulfate chain at position S583, which distinguishes it from the Sulf-1 enzyme (El Masri et al., 2022). The activity of Sulf-2 is regulated by the chondroitin sulfate (El Masri et al., 2022; Vivès et al., 2014) and by its N-glycosylation (Benicky et al., 2023). It is also known that the catalytic activity is sensitive to ionic influences and buffer composition (Benicky et al., 2023; Morimoto-Tomita et al., 2002). However, virtually nothing is known about regulation of the enzymatic activity by interaction with other proteins.

We anticipated that the interactions of the Sulf-2 enzyme with other protein in the extracellular space regulate its function. To test our hypothesis, we performed monoclonal antibody (mAb) affinity purification of the native Sulf-2 and its interacting partners from the secretome of head and neck squamous cell carcinoma (HNSCC) cell lines. We identified and validated highly specific binding of Sulf-2 to galectin-3-binding protein (Gene: LGALS3BP, UniProtKB Q08380). Furthermore, using *in vitro* enzymatic and cell culture assays, we showed that the interaction inhibits Sulf-2 enzymatic activity and modifies invasion of cancer cells into matrigel.

## Results

### Identification of Sulf-2 interacting proteins

HNSCC cell lines SCC35 and CAL33 (human squamous cell carcinoma cell lines derived from hypopharynx and oral cavity tumors, respectively) were grown in 75 cm^2^ flasks as described (Mukherjee et al., 2023) and serum-free conditioned media were collected for 24h. The cleared supernatant (secretome) was used as the starting material for immuno-affinity purification (IP) of the Sulf-2 protein in complex with its interacting partners (Supplementary Figure 1). Sulf-2 specific monoclonal antibodies (Singer et al., 2015) bound to magnetic beads were used to pulldown the target protein, and the co-purified proteins were identified by LC-MS/MS analysis. The identified proteins including proteins detected in parallel negative control are listed in Supplementary Table 1. As expected, the Sulf-2 protein was only identified in specific mAb pulldowns but not in antibody isotype negative control. Additionally, we identified several proteins in the pulldown samples that were not identified in negative control and do not belong to known common contaminants (Table 1). Of the candidates, Sulf-2 and LG3BP were consistently identified in replicate experiments, including IP samples prepared under high stringency wash with 200 mM NaCl (results not shown). From these results we infer that the Sulf-2 protein interacts with LG3BP *in vivo*.

**Table 1.** Candidate interacting partners of the Sulf-2 protein identified in anti-Sulf-2 mAb affinity pull-downs.

| Accession | Description | Exp. q-value:<br>Combined | Sum<br>PEP<br>Score | Coverage<br>[%] | #<br>Peptides | #<br>PSMs |
| --- | --- | --- | --- | --- | --- | --- |
| Q08380 | Galectin-3-binding protein | 0 | 56.372 | 41 | 18 | 53 |
| Q8IWU5 | Extracellular sulfatase Sulf-2 | 0 | 26.877 | 17 | 13 | 26 |
| P05023 | Sodium/potassium-transporting ATPase subunit alpha-1 | 0 | 25.134 | 16 | 13 | 26 |
| Q8WWM7 | Ataxin-2-like protein | 0 | 18.231 | 12 | 11 | 16 |
| Q7Z417 | Nuclear fragile X mental retardation-interacting protein 2 | 0 | 24.334 | 21 | 9 | 14 |
| Q9UN86 | Ras GTPase-activating protein-binding protein 2 | 0 | 25.357 | 21 | 9 | 31 |
| P11021 | Endoplasmic reticulum chaperone BiP | 0 | 12.428 | 16 | 7 | 11 |
| P29692 | Elongation factor 1-delta | 0 | 10.734 | 22 | 4 | 8 |
| Q9BTT6 | Leucine-rich repeat-containing protein 1 | 0 | 6.083 | 6 | 3 | 4 |
| Q5TC12 | ATP synthase mitochondrial F1 complex assembly factor 1 | 0 | 4.611 | 7 | 2 | 4 |
| Q08554 | Desmocollin-1 | 0 | 4.563 | 3 | 2 | 3 |
| Q99729 | Heterogeneous nuclear ribonucleoprotein A/B | 0 | 4.199 | 7 | 2 | 3 |

### Validation of the Sulf-2 and LG3BP interaction

We generated a recombinant Sulf-2 (rSulf-2) protein in HEK293T cells using lentiviral transduction, and performed affinity purification of the Sulf-2 protein from the serum free media of the transduced cell line using Sulf-2 mAb. Reciprocally, LG3BP protein pulldown from this sample was performed using anti-LG3BP monoclonal antibody. MS analysis identified Sulf-2 and LG3BP proteins in both pulldowns (Figure 1A and Supplementary Table 2). We also verified presence of these two proteins in pulldown sample, and not in negative control samples, by Western blot analysis (Figure 1B).

**Figure 1.**
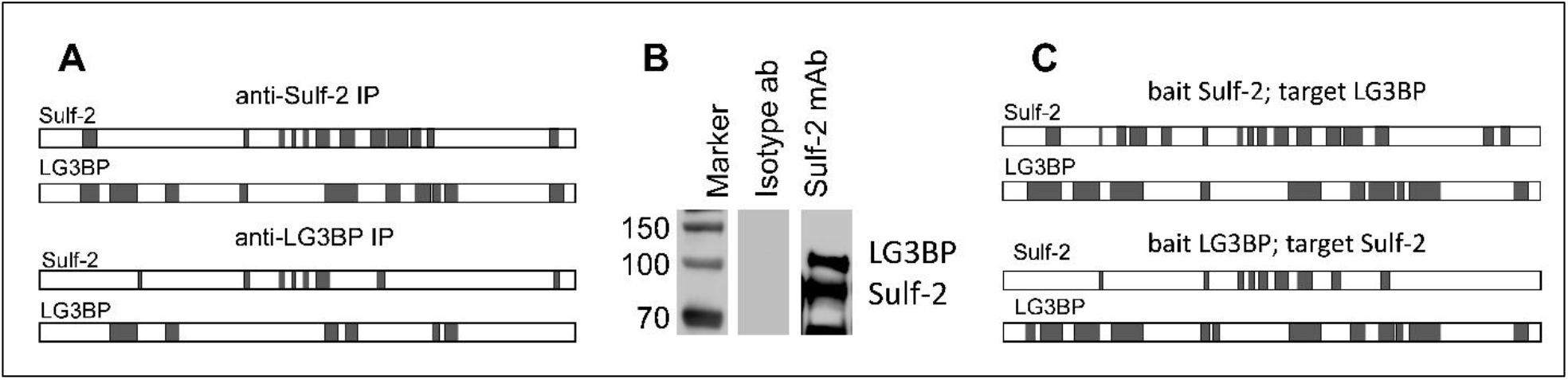
Identification of proteins in affinity pulldown samples. (A) LC-MS/MS analysis identified Sulf-2 and LG3BP proteins by multiple peptide matches (shaded region) in IP samples from HEK293T culture media using anti-Sulf-2 or LG3BP mAbs as indicated. (B) Western blot analysis of the IP samples prepared from HEK293T culture using anti Sulf-2 mAb, and isotype antibody as negative control. Blot was probed with anti-Sulf-2 and LG3BP mAbs. (C) Sulf-2 and LG3BP were identified by LC-MS/MS (matched peptide indicated by shaded region) in reciprocal co-IP samples using purified recombinant proteins, the bait and target as noted.

In addition, we verified the direct interaction of Sulf-2 and LG3BP *in vitro* by reciprocal co-immunoprecipitation (co-IP) experiments using purified recombinant proteins (rSulf-2, rLG3BP). The rSulf-2 protein bound to the beads (bait) was incubated with rLG3BP (target) and vice-versa in parallel (see methods). Following extensive washes, Sulf-2 and LG3BP were both identified by MS analysis in both co-IP experiments but not in negative controls (Figure 1C). These results unequivocally confirm the binding of Sulf-2 and LG3BP.

### Effect of chondroitin sulfate on Sulf-2 interaction with LG3BP

We interrogated if the chondroitin sulfate (CS) covalently attached to Sulf-2 at S583 influences its interaction with LG3BP. To assess this, we first treated the rSulf-2 media with chodroitinase-ABC, followed by affinity pulldown using anti-LG3BP mAb. A control sample was prepared in parallel under identical conditions except for the chodroitinase addition. The relative abundance of the Sulf-2 and LG3BP proteins in IP samples was determined by targeted LC-MS/MS-PRM of a Sulf-2 tryptic peptide AEYQTAcEQLGQK (z +2, *m/z* 763.3512) and LG3BP tryptic peptide YSSDYFQAPSDYR (z +2, *m/z* 799.8415) as detailed in methods. The ratio of the peak areas showed a 3-fold increase in LG3BP binding to Sulf-2 upon the CS-side chain removal (Figure 2). The results suggest that the CS side-chain influences the interaction, either directly or by modifying the Sulf-2 structure.

**Figure 2.**
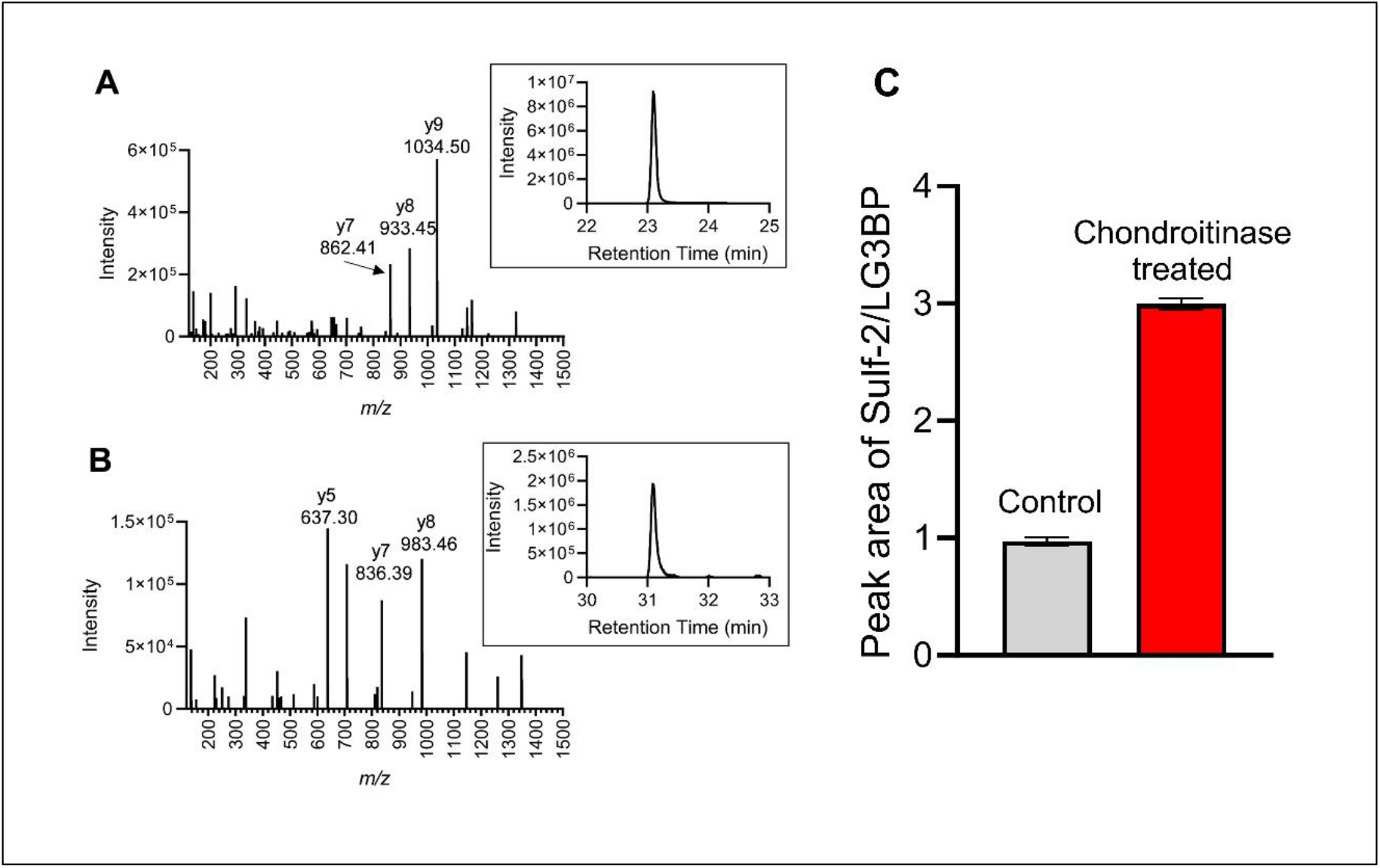
LC-MS/MS PRM analysis of Sulf-2 and LG3BP in LG3BP mAb pulldown of chondrotinase treated and control samples. Product ion spectra and extracted peaks using sum of indicated product ions of the following peptides: (A) Sulf-2 target peptide AEYQTAcEQLGQK (*z* +2, *m/z* 763.3512); and (B) LG3BP target peptide YSSDYFQAPSDYR (*z* +2, *m/z* 799.8415). (C) Graphical representation of the average ratio of peak areas (triplicate runs) of Sulf-2 / LG3BP peptides, showing 3-fold relative increase of Sulf-2 in the chondroitinase treated pulldown sample.

To confirm the impact of the CS on binding, we prepared a S583A mutant of rSulf-2 in a HEK293F expression system as described previously (Benicky et al., 2023). The lack of CS on the S583A mutant rSulf-2 was verified by native gel electrophoresis followed by western blot analysis (Figure 3A); this is in contrast to the wild-type protein which is almost completely decorated with the CS side-chain. However, a significant proportion of lentiviral expressed rSulf-2 (HEK293T cells) lacks the CS side chain. Subsequently, we determined the relative abundance LG3BP in the complex with the S583A Sulf-2 by the LC-MS/MS-PRM assay in pulldown samples prepared with anti Sulf-2 mAb. The relative abundance of LG3BP was highest in the Sulf-2 CS mutant, followed by wild type Sulf-2 expressed in the HEK293T lentiviral sample (which has substantial proportion of Sulf-2 lacking the CS), and lowest in wild type Sulf-2 expressed in the HEK293Fcell (nearly completely CS-modified) (Figure 3B). The results show an inverse correlation of the binding to the CS content of the Sulf-2 preps. The CS chain clearly impacts the binding of LG3BP to the Sulf-2 protein and may impact their interaction *in vivo*.

**Figure 3.**
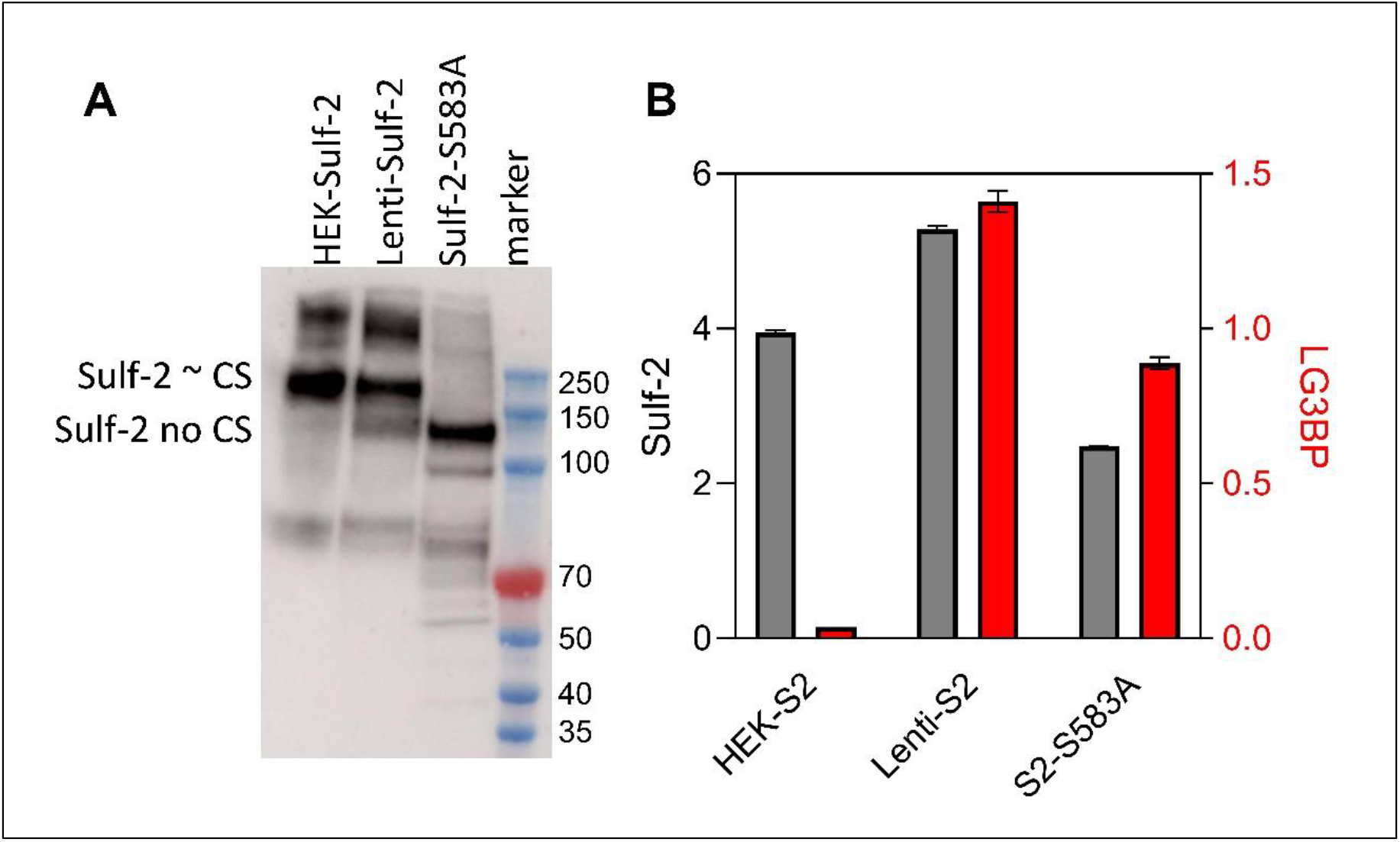
Effect of the Sulf-2 CS chain on LG3BP interaction. A. Western blot analysis (non-reduced sample) showing Sulf-2 mostly chondroitinated in HEK, non-chondrotinated in S583A mutant and mixture of chondroitinated and non-chondroitinated in lenti Sulf-2 media. B. LC-MS/MS-PRM quantification of Sulf-2 (grey bars) and gal3bp (red bars) in Sulf-2 mAb pulldown samples shows relative increase of gal3bp binding in samples containing more non-chondroitinated Sulf-2.

### Effect of LG3BP on Sulf-2 enzymatic activity

Sulf-2 enzymatic activity was measured in the presence and absence of LG3BP using a 2S2-6S4 synthetic oligosaccharide substrate as described recently (Benicky et al., 2023). Purified rSulf-2 was incubated with varying concentrations of purified rLG3BP followed by the addition of a fixed concentration of the 2S2-6S4 substrate. The desulfation of the substrate by Sulf-2 was inhibited by LG3BP in a dose dependent manner (Figure 4). At a 4h time point, the Sulf-2 activity was reduced by approximately 50% at a LG3BP concentration 8 μg/ml which corresponds to a 40x excess over the Sulf-2 enzyme in the reaction. The result confirms a direct interaction of the Sulf-2 and LG3BP proteins and shows that the interaction has functional consequences which could impact the Sulf-2 activity in model systems and *in vivo*.

**Figure 4.**
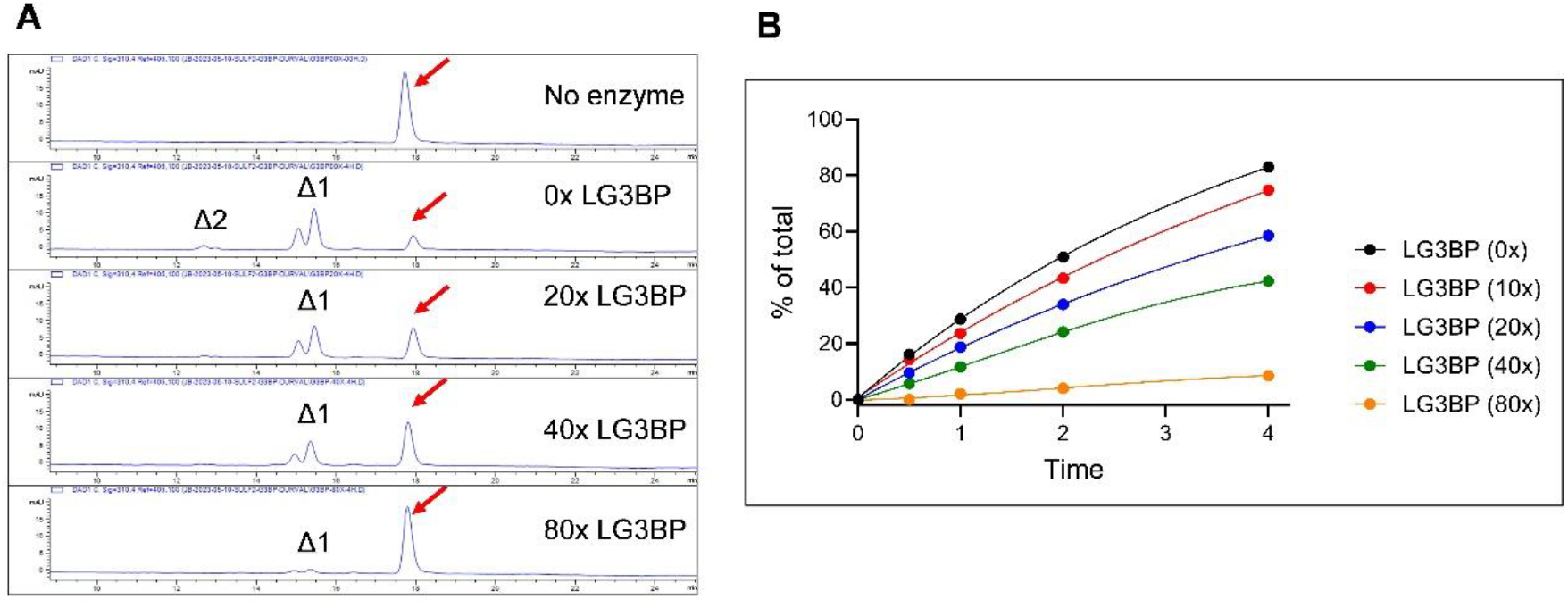
Measurement of the Sulf-2 enzymatic activity with LG3BP added at increasing concentrations. (A) Representative chromatograms showing the substrate 2S2-6S4 (indicated by red arrow) and product (Δ1, Δ2) peaks where the products with 1 or 2 sulfates removed decrease with increasing LG3BP additions. (B) Percent of converted substrate over a 4h time course with fixed concentration of Sulf-2 and increasing concentrations of LG3BP expressed as fold-excess over the Sulf-2 concentration.

### Inhibition of cancer cell invasion by LG3BP in a spheroid model

We recently reported a Sulf-2 dependent spheroid invasion model consisting of a co-culture of SCC35 cancer cells with a primary cancer associated fibroblast (HNCAF37) in Matrigel (Mukherjee et al., 2023). We have shown that Sulf-2 KO SCC35 cells invade Matrigel in this system significantly less than the wild type SCC35 cells. In this study, we used the same model to assess how an exogenous addition of LG3BP to the spheroid model affects cancer cell invasion (Figure 5). The control spheroids significantly invaded Matrigel by day 5, as expected. We observed significant reduction in the invasion phenotypes on day 5 in the LG3BP treated spheroids. Compared to the no treatment control (NTC), area of the spheroids was reduced to 66%, inverse circularity to 19%, protrusion to 49%, and endpoints to 36% with the addition of 100 ng LG3BP. No significant differences were observed between day 5 spheroids treated with 100 ng or 1μg of LG3BP, indicating saturation of the binding already at 100 ng. Subsequently, we used SCC35 Sulf-2 KO cell in the spheroid assay and we observed a reduced influence of the LG3BP. The NTC Sulf-2 KO spheroids invaded Matrigel less efficiently than the NTC Sulf-2 wild type spheroids (reduction of area to 35%, inverse circularity to 24%, protrusion to 51%, and endpoints to 67%), as expected, but the addition of LG3BP did not significantly alter the outcome in the Sulf-2 KO spheroids. Thus, addition of LG3BP affected invasion of the wild type SCC35 cells into Matrigel in a Sulf-2 dependent manner. This confirms that the inhibition of Sulf-2 enzymatic activity could have functional consequences *in vivo*.

**Figure 5.**
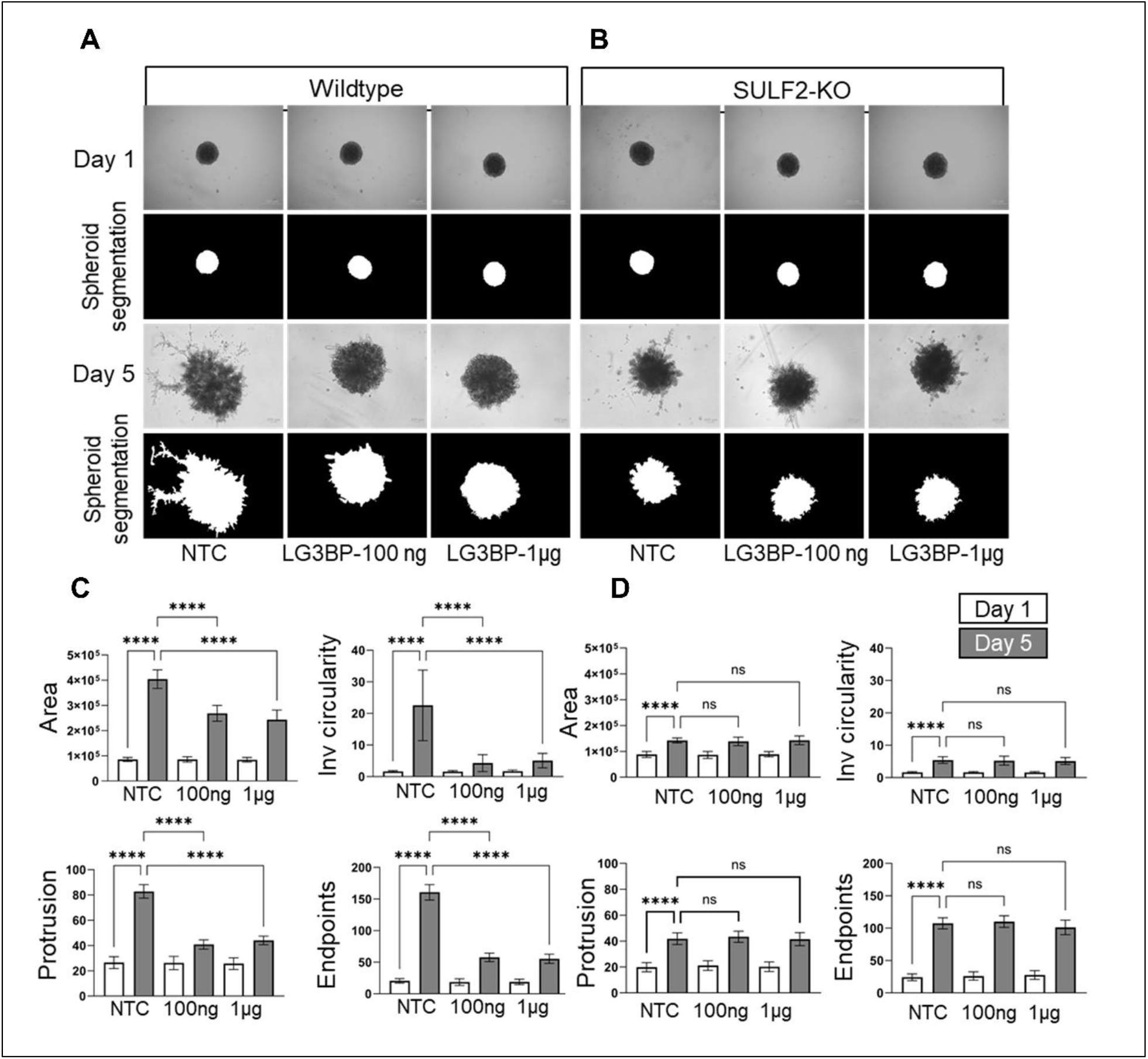
Effect of LG3BP addition on the invasion of SCC35 spheroids in Matrigel. Upper left panel (A) shows assays using WT SCC35 cells co-cultured with HNCAF61137 cells, right panel (B) shows SCC35 SULF2-KO (knock out) cells co-cultured with HNCAF61137 cells. Representative bright field images with phase contrast (top row) and masked greyscale images generated with INSIDIA (bottom row) of the co-culture spheroids grown in Matrigel and imaged on day 1 and day 5. Scale bar 200 μm. Lower panel graphical representation of spheroid invasion (day 1 open box, day 5 shaded box) measured without LG3BP (NTC) and with two concentrations of LG3BP (100 ng and 1 μg) as indicated in the following co-cultures: C. SCC35 wild type spheroids; D. SCC3 SULF2 KO spheroids. Results are reported as total area, inverse circularity, protrusions, and endpoints. Bars indicate mean ± SD with statistical significance evaluated using two-way ANOVA with post hoc Tukey’s test. P-values <0.05 are considered significant, p-values <0.0001 are represented by ^****^.

## Discussion

Heparan-6-*O*-endosulfatases Sulf-1 and Sulf-2 are secreted extracellular enzymes overexpressed in many types of cancer (Flowers et al., 2016; Hur et al., 2012; Lai et al., 2008; Lee et al., 2017; Yang et al., 2022, 2021; Zhu et al., 2016). The Sulf-1 protein is upregulated in the tumor tissues, but in general is not detectable in cancer cell lines. Sulf-2 is secreted in the tumor tissues primarily by cancer cells while Sulf-1 is supplied to the tumor microenvironment by CAF cells. The Sulf-2 is considered oncogenic (Rosen and Lemjabbar-Alaoui, 2010) and we have shown that Sulf-2 is associated with poor survival outcomes in HNSCC (Yang et al., 2022, 2021). Even though Sulf-1 and Sulf-2 share many common features and *in vitro* enzymatic activity, only Sulf-2 is a proteoglycan modified by a CS chain reported to affect its enzymatic activity (El Masri et al., 2022; Vivès et al., 2014). However, it remains unknown what other factors regulate activity of the Sulf enzymes *in vivo*. We suspected that the activity of Sulf-2 depends on interacting regulatory proteins, but virtually nothing is known about its activity-modifying interactions. In this study, we therefore searched for Sulf-2 interacting proteins that modify its enzymatic activity.

Because Sulf-2 is a secreted glycoprotein, it is expected that the interacting proteins will be found in the extracellular environment. We therefore focused on the interacting partners in the secretome of HNSCC cancer cell lines expressing the Sulf-2 protein. We used an affinity pulldown approach using specific anti-Sulf-2 mAbs and mass spectrometry to identify its interacting partners. Subsequently, we validated the interaction of our top candidate LG3BP, and documented its impact on the activity of the Sulf-2 enzyme. The interaction of LG3BP with proteins of various biological function and cellular location has been reported in protein-protein interaction studies (Bouwmeester et al., 2004; Haenig et al., 2020; Hubel et al., 2019). A recent study (Siegel et al., 2022) showed that Sulf-2 binds TNFR1 in TNF-α-stimulated primary human rheumatoid arthritis synovial fibroblasts (RASFs). Additional candidate interactors (Amyloid-beta precursor protein, Dickkopf-related protein 3, Neutrophil defensin 1, Defensin alpha 5) of Sulf-2 were observed in a high-throughput interactome study (Huttlin et al., 2017). However, these candidates were not verified and their impact on Sulf-2 activity was not examined. To our knowledge the interaction of Sulf-2 with LG3BP has not been reported, and we are first to report this interaction in a targeted validated study.

We identified LG3BP in Sulf-2 pulldown from the secretomes of two different HNSCC cell lines under high stringency washes. The association of the two proteins was further verified by complementary pull down of Sulf-2 and LG3BP with their respective mAbs from the conditioned media of cell lines over expressing a recombinant Sulf-2 protein (Figure 1). It is possible that the LG3BP associates with Sulf-2 in a complex with other proteins because we identified other candidate proteins in the pulldown samples after subtraction of a negative pull down control and common contaminants (Table 1). However, LG3BP interacts with Sulf-2 directly, independent of the other proteins (see below). We therefore focused on characterizing the Sulf-2 interaction with LG3BP and evaluated its impact on Sulf-2 enzymatic activity.

To confirm that the Sulf-2 and LG3BP bind directly, we performed reciprocal co-IP experiments using purified recombinant proteins (Figure 1C). Our results show clearly a direct interaction of the two proteins and we infer that they likely directly interact *in vivo*. However, it is interesting to note that the interaction depends on the CS modification of Sulf-2. Treatment of Sulf-2 with chondroitinase ABC increases efficiency of the LG3BP pulldown (Figure 2). In addition, we observed in pulldown experiments with proteins from our recombinant cell cultures that LG3BP binds weaker to the chondroitinated Sulf-2 (Figure 3). We showed by targeted MS assays that the relative abundance of LG3BP in the pulldown samples is inversely related to the chondroitination of Sulf-2. To validate this observation, we generated a mutant Sulf-2 that lacks the CS-chain and we find that it pulls down LG3BP more efficiently (Figure 3). Thus, the CS-side chain of SULF2 may be an important modifier of this interaction and the degree of chondroitination of the Sulf-2 enzyme in different cells and tissues needs to be further examined together with the structure of the CS chain.

LG3BP is a relatively abundant secreted protein ubiquitously present in human tissues and detectable in serum. It is a multi-function glycosylated protein, an abundant component of cancer‐derived EVs, and a key regulator of cancer-stroma interactions (reviewed in Capone et al., 2021). It has been described as a biomarker and as a therapeutic target (Dufrusine et al., 2023). Overall, it is a substantially more abundant protein than the Sulf-2 enzyme. It is also nearly 100 times more abundant in serum than galectin 3, its well-recognized binding partner (Iurisci et al., 2000). LG3BP is detected in human serum at concentrations of approximately 3 μg/ml, is further elevated in the sera of cancer patients, and could have even higher titers in the tumor microenvironment (Iurisci et al., 2000; Nielsen et al., 2017). It is also an abundant protein in the secretomes of all our HNSCC cancer and CAF cell lines which further supports our assumption that it interacts with Sulf-2 *in vivo*.

We assessed the functional significance of the Sulf-2 interaction with LG3BP *in vitro* by an enzymatic assay (Figure 4), Our results show that LG3BP inhibited Sulf-2 activity in a concentration dependent manner at concentrations and inhibitor/enzyme ratio’s plausible *in vivo*. The formation of Sulf-2 LG3BP complex in biological samples might be affected by additional factors not observed when we compare the interaction of two purified recombinant proteins; we have already shown the influence of the CS chain which was not yet evaluated sufficiently *in vivo*. However, it is very plausible there two proteins have direct interaction *in vivo*. Similarly, the impact of LG3BP on Sulf-2 activity *in vivo* will need further evaluation even though our results show that the interaction is a plausible regulator of the Sulf-2 enzymatic activity. This is further supported by our spheroid co-culture models. We have shown that the addition of purified recombinant LG3BP to the spheroid culture media inhibited invasion of the SCC35 cells into Matrigel. In addition, invasion of the Sulf-2 KO SCC35 cells is not affected by the addition of LG3BP which is similar to a previously observed effect of a Sulf-2 inhibitor in this model system (Mukherjee et al., 2023). Thus, we conclude that LG3BP directly interacts with Sulf-2 and inhibits the Sulf-2 dependent cancer cell invasion.

In summary, we have shown that the Sulf-2 enzyme interacts directly with LG3BP. This interaction inhibits the Sulf-2 enzymatic activity and is modified by the CS modification of Sulf-2. We propose that this interaction represents a plausible *in vivo* regulatory mechanism of the Sulf-2 editing of heparan 6-*O*-sulfation. The results strongly support the need for further evaluation of this regulatory interaction in relevant physiological and pathologic conditions.

## Materials and Methods

### Materials

Amicon Ultra-15 (Millipore), Dynabeads™ M-280 sheep anti-Mouse IgG (Invitrogen), Anti-Sulf-2 monoclonal antibodies (QED Bioscience), Anti-LG3BP monoclonal antibody (Proteintech), Mouse IgG1 k Isotype control (eBioscience), BSA (NEB), Recombinant Sulf-2 protein (produced in house), Recombinant human galectin 3 binding protein (R&D Systems, and produced in-house), QuikChange Lightning Site-Directed Mutagenesis Kit (Agilent Technologies), 25 kDa polyethylenimine (Polysciences), Custom heavy isotope labeled Sulf-2 peptide (Biosynth), Trypsin/Lys-C protease (Thermo Scientific), Empore C18 cartridge (CDS Analytical), Nano-flow C18 columns, and other reagents and chemicals were obtained from Thermo Scientific.

### Cell culture and secretome collection

Two HNSCC cell lines, SCC35 and CAL33, were cultured as described previously (Mukherjee et al., 2023). Briefly, the cells were grown to 70% confluency in DMEM containing 10% FBS, following which they were washed with and incubated in pre-warmed (37°C) serum free conditioned media for 24h at 37°C. The conditioned media were collected and non-ionic detergent Tween-20 was added to a final concentration of 0.01%. Cells and any cell debris were removed by two-step centrifugations, 800 g for 10 min at 4°C, followed by 4,000 g for 10 min at 4°C. The supernatant was concentrated to 50x using Amicon Ultra-15 Centrifugal Filters (30kDa cutoff), and further centrifuged at 16,200 g for 15 min to remove any aggregates. The cleared concentrated secretome was used in our immunoprecipitation assays.

### Production of recombinant Sulf-2 and LG3BP using a lentiviral expression system

SULF2 ORF including C-terminal Myc-His tag was subcloned from its source vector (Addgene # 13003) to lentiviral transfer vector pHR-CMV-TetO2_3C-Twin-Strep (Addgene # 113883) (Elegheert et al., 2018). Galectin-3-binding protein ORF was subcloned from its source vector (Origene # RC204918) into bicistronic lentiviral transfer vector pHR-CMV-TetO2 3C-TwinStrep-IRES-EmGFP vector (Addgene, # 113884) modified to introduce 6xHis tag downstream of a cleavable C-terminal TwinStrep tag. The above transfer vectors were co-transfected with envelope (pMD2.G, Addgene) and packaging (psPAX2, Addgene) plasmids to Lenti-X HEK293T cells (Takara) to generate lentiviral particles for transduction of the target cells as described in detail previously (Elegheert et al., 2018). The His-tagged Sulf-2 (Q8IWU5) and LG3BP (Q08380) proteins were purified using nickel affinity column as described earlier (Benicky et al., 2023). SDS-PAGE protein profile of purified LG3BP sample is shown in Supplementary Figure 2.

### Generation of mutant Sulf-2 protein without chondroitin sulfate

We described previously (Benicky et al., 2023) production of a recombinant Sulf-2 protein using a stably transfect HEK293F cell line. We used the same expression system to generate a cell line expressing a Sulf-2 protein with a Ser583Ala (S583A) point mutation, which eliminates the chondroitin sulfate (CS) attachment site (position) from the expression vector pcDNA3.1/Myc-His(-)-HSulf-2 bearing human Sulf-2 cDNA (a gift from Steven Rosen, Addgene plasmid # 13004) (Morimoto-Tomita et al., 2002). The Sulf-2 S583A mutation was generated using mutagenic primers 5’-GGCCTCCAGTGCCAGCGAAGTCCCCACCAT-3’ (sense) and 5’-ATGGTGGGGACTTCGCTGGCACTGGAGGCC-3’, and site-directed mutagenesis kit, according to the manufacturer’s instructions. The construct was transfected into HEK293F cells using linear 25 kDa polyethylenimine at nitrogen-to-phosphate (N/P) ratio 20. Stable transfectants were generated by antibiotic resistance selection using 500 μg/ml G418.

### Immunoprecipitation of native Sulf-2 and interacting proteins

The native Sulf-2 protein was immunoprecipitated from the secretome of the Cal33 and SCC35 HNSCC cell lines using anti-Sulf-2 monoclonal antibodies (mAbs 5D5 and 5C12). The following steps were carried out at 4°C. The mAbs (5 μg each diluted in 500 μl of IP buffer: 1x PBS, 0.1% Tween 20) were incubated with 100 μl of anti-mouse IgG coated magnetic beads (Dynabeads M-280) and 10 μg BSA for 1 h with rotation. Beads incubated with mouse IgG isotype served as negative control. The beads were washed thrice with IP buffer (1 ml each step) and the antibody bound beads were incubated with the secretome sample (500 μl of the concentrated SCC35 or CAL33 in parallel plus 500 μl IP buffer) for 1h with rotation. Unbound materials were removed by washing the beads three times with the IP buffer (1ml each step, 5 min each incubation, with rotation). The beads were further washed thrice with 1x PBS to remove residual detergent from the sample (for ease of downstream mass spectrometry analysis). The bound proteins were eluted by incubating the beads in 10 mM Glycine pH 2.6, and neutralized with Tris buffer. The eluates were processed for mass spectrometry analysis as described below.

### Immunoprecipitation of recombinant Sulf-2 and interacting protein from conditioned media

We collected cleared conditioned media from HEK293F or HEK293T cells expressing recombinant Sulf-2 protein, and HEK293F cells expressing S583A-Sulf-2 mutant protein. The media was concentrated 5x using a 30 kDa cut-off membrane, and cleared by centrifugation as described above. The Sulf-2 and its interacting protein(s) were pulled down from these samples using anti-Sulf-2 monoclonal antibodies as above. In parallel, LG3BP and associated proteins were pulled down using anti-LG3BP mAb, the IP steps were carried out as above using 5 μg mAb. The bound proteins were eluted with 10 mM glycine as above, and used for LC-MS/MS or western blot analysis.

### Reciprocal Co-IP analysis

Co-immunoprecipitation experiments were performed using purified recombinant proteins to validate their interaction. Briefly, anti-mouse magnetic beads (50 μl suspension) were incubated with anti-LG3BP monoclonal antibody in IP buffer and washed as above. Then the beads were incubated with purified recombinant LG3BP (source R&D Systems) and washed thrice with IP buffer. The LG3BP bound beads were incubated with purified recombinant Sulf-2 proteins, and washed with IP buffer to remove unbound proteins. Simultaneously, reciprocal pulldown experiment was performed where the beads were first incubated with Sulf-2 mAb, then with purified Sulf-2 protein, followed by LG3BP protein. The bound proteins were eluted as above, and analyzed by mass spectrometry as described below.

### SDS-PAGE and Western Blot analysis

The proteins in immunoprecipitated sample were denatured in SDS-PAGE loading buffer with or without DTT, and were separated by SDS-PAGE using NuPAGE 4–12% polyacrylamide gels.

Western blot was performed with Sulf-2 (1:2500 dilution) and LG3BP mAbs (1:5000 dilution).

### LC-MS/MS analysis

The proteins in immunoprecipitated samples were reduced/alkylated by treatment with TCEP (10 mM) and iodoacetamide (6 mM), followed by in solution proteolytic digestion in Barocycler with Lys-C/Trypsin (200 ng in 100 μl volume). The resulting peptides were desalted using a C18 cartridge and analyzed by mass spectrometry using a Dionex RSLCnano – Orbitrap fusion lumos LC-MS platform configured with Acclaim PepMap 100 trap column (75 μm x 2 cm, C18, 3 μm, 100A), and PepMap RSLC C18 EASY-spray analytical column (75 μm x 15 cm, 3 μm, 100A). The peptides were separated using a 60-min acetonitrile gradient at a flow rate of 0.3 μl/min; solvent A (0.1% formic acid in water), solvent B (0.1% formic acid in acetonitrile), 0-5 min peptide load to trap column at 2% B (valve switched at 5 min to direct the flow to analytical column), 5-35 min 5-25% B, and 35-45 min 25-90% B, 45-53 min 90%B, 53-60 min 2% B. The Orbitrap MS parameters were set at nanospray voltage of 2.0 kV, capillary temperature 275°C, MS1 scan range m/z 400–1800, resolution 120,000, RF lens 60%, and MS2 intensity threshold 2.0 × 10^4^, included charge states 2–7, and dynamic exclusion for 15s. Data dependent HCD MS/MS spectra were collected at 7,500 resolution with 30% collision energy. The acquired mass spectrometry data were searched against human uniport database using Proteome Discoverer Sequest HT search engine, and filtered with Percolator algorithm.

### Targeted LC-MS/MS-PRM analysis

The relative quantities of Sulf-2 and LG3BP in IP samples were determined by targeted mass spectrometry. Briefly, the IP samples prepared in parallel from HEK cell conditioned media (recombinant Sulf-2 and mutant Sulf-2) were digested and spiked with 5 femtomole/μl of a heavy (C^13^, N^15^) labeled Sulf-2 internal peptide AEYQTACEQLGQK*. Targeted LC-MS/MS-PRM assay was set to collect the product ions of native Sulf-2 peptide (*m/z* 763.3512, *z* 2), heavy Sulf-2 peptide (*m/z* 767.3583, *z* 2), and LG3BP peptide YSSDYFQAPSDYR (*m/z* 799.8415, *z* 2). The LC parameters were as above, the HCD product ion spectra of the selected precursors were collected at 1.6 *m/z* isolation window, 30% collision energy and 15k orbitrap resolution.

Xcalibur Quan Browser software was used to extract the peak area of the peptides by sum of 3 product ions; AEYQTACEQLGQK (y7 862.4087, y8 933.4458, y9 1034.4935), AEYQTACEQLGQK* (y7 870.4229, y8 941.4600, y9 1042.5077), and YSSDYFQAPSDYR (y5 637.2940, y7 836.3897, y8 983.4581). For each sample the assay was performed in triplicate and the peak areas of the peptides were normalized to the spiked heavy peptide; and relative abundance of Sulf-2 to LG3BP in the pulldown sample was assigned based on the peak area of the target peptides.

### Sulf-2 enzymatic activity assay

Effect of LG3BP on Sulf-2 enzymatic activity was measured using 2S2-6S4 substrate, a recently described HPLC-UV assay (Benicky et al., 2023). Briefly, LG3BP (R&D Systems) at a 2-fold dilution series (concentration range of 2-16 μg/ml) was preincubated with a fixed amount of Sulf-2 (0.4 μg/ml) for 30 min before addition of 100 μM 2S2-6S4 substrate. BSA was added in place of LG3BP (at highest equimolar concentration) to the Sulf-2 as a control. The reaction was carried out in a 50 μl volume, and aliquots were collected at 0, 0.5, 1, 2, and 4 h. The substrate and product(s) were quantified by ion exchange HPLC with UV detection.

### Spheroid assay

The spheroid assays using SCC35 and HNCAF61137 cells were performed as described (Mukherjee et al., 2023). Briefly, the spheroids were formed by co-culturing the cells for 1 day in ultra-low attachment 96-well round bottom plates and were embedded in 50 μl of growth factor reduced Matrigel. Then 100 μl medium with / without LG3BP (100 ng/well or 1 μg/well; 6 replicates per concentration) was gently added on top of the Matrigel in each well. Spheroids were cultured for 5 days and imaged on day 1 and day 5 using Olympus IX71 inverted microscope. Digital images were analyzed for spheroid area, inverse circularity, protrusion, and endpoint measurements using INSIDIA 2.0 software (Moriconi et al., 2017; Perini et al., 2022). GraphPad Prism Software (Version 10.1.1) was used to perform statistical analysis and data visualization. Mean and SD was calculated, and two-way ANOVA with Tukey was performed to determine statistical significance between data groups.

## Funding

This work was supported in part by a National Institutes of Health grants R01CA238455 to RG. The content is solely the responsibility of the authors and does not necessarily represent the official views of the National Institutes of Health. This work was also partly supported by the CAS (RVO: 86652036) and the National Institute for Cancer Research (Programme EXCELES, ID Project No. LX22NPO5102) to CB.

## Acknowledgements

We thank Laurie Ailles, University of Toronto for providing cell lines and advice on spheroid assays. Further support was provided by the Office of The Director, National Institutes of Health under Award Number S10OD023557 supporting the operation of the Clinical and Translational Glycoscience Research Center, Georgetown University.

## Author contributions

Conceptualization, AP, JB, and RG; methodology and formal analysis AP, JB, RA, PM and ZN; investigation, AP, JB, and PM; resources, CB, RG; data curation, AP, JB, PM; writing - original draft preparation, AP, JB, and PM; review and editing, AP, JB, CB, and RG; supervision; project administration, RG; funding acquisition, RG. All authors have read and agreed to the published version of the manuscript.

## Competing interests

The authors declare no competing interests.

## Notes

### Competing Interest Statement

The authors have declared no competing interest.

