## Supplementary Figures for "Galectin-3-binding protein inhibits extracellular heparan 6-*O*-endosulfatse Sulf-2"

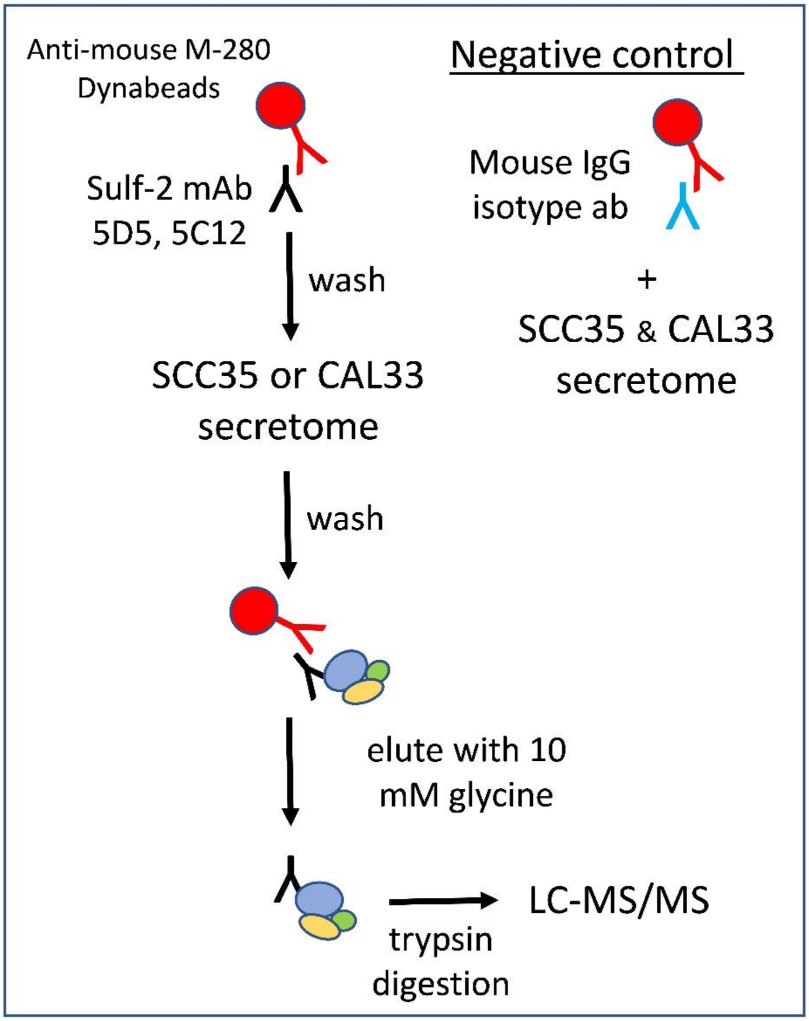


**Supplementary Figure 1**. Schematics of the affinity pull-down method of Sulf-2 and associated proteins using monoclonal antibodies.

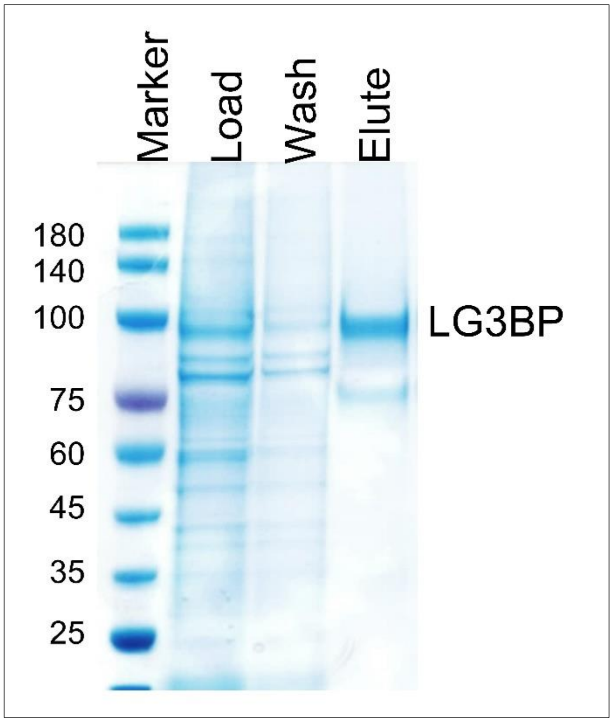


**Supplementary Figure 2**. Coomassie stained SDS-PAGE of $Ni^{2+}$ affinity purified LG3BP (lane-Elute). Protein profile of starting conditioned media (Load) and from washing step (Wash) is shown.
